# The SpyBLI cell-free pipeline for the rapid quantification of binding kinetics from crude samples

**DOI:** 10.1101/2025.03.06.641810

**Authors:** Olga Predeina, Misha Atkinson, Oliver Wissett, Montader Ali, Cristina Visentin, Stefano Ricagno, Anthony H. Keeble, Mark R. Howarth, Pietro Sormanni

## Abstract

Accurate measurements of binding kinetics, encompassing equilibrium dissociation constant (K_D_), association rate (*k*_on_), and dissociation rate (*k*_o,_), are critical for the development and optimisation of high-affinity binding proteins. However, such measurements require highly purified material and precise ligand immobilisation, limiting the number of binders that can be characterised within a reasonable timescale and budget. Here, we present the SpyBLI method, a rapid and cost-effective biolayer interferometry (BLI) pipeline that leverages the SpyCatcher003–SpyTag003 covalent association, eliminating the need for both binder purification and concentration determination. This approach allows for accurate binding-kinetic measurements to be performed directly from crude mammalian-cell supernatants or cell-free expression mixtures. We also introduce a linear gene fragment design that enables reliable expression in cell-free systems without any PCR or cloning steps, allowing binding kinetics data to be collected in under 24 hours from receiving inexpensive DNA fragments, with minimal hands-on time. We demonstrate the method’s broad applicability using a range of nanobodies and single-chain antibody variable fragments (scFvs), with affinity values spanning six orders of magnitude. By minimising sample preparation and employing highly controlled, ordered sensor immobilisation, our workflow delivers reliable kinetic measurements from crude mixtures without sacrificing precision. We expect that the opportunity to carry out rapid and accurate binding measurements in good throughput should prove especially valuable for binder engineering, the screening of next-generation sequencing–derived libraries, and computational protein design, where large numbers of potential binders for the same target must be rapidly and accurately characterised to enable iterative refinement and candidate selection.

## Introduction

Developing and characterising high-affinity binding proteins, such as antibodies, critically depends on precise measurements of their interactions with target antigens^1^. Accurate quantification of these interactions informs the selection of candidates through discovery and optimisation, and ultimately influences efficacy and specificity of the final product. Traditional methods like enzyme-linked immunosorbent assays (ELISAs) estimate binding affinity through proxies such as the half-maximal effective concentration (EC_50_). While useful, these measurements offer limited insight into full binding kinetics – including the equilibrium dissociation constant (K_D_), association rate (*k*_on_), and dissociation rate (*k*_o,_) – which are essential for a comprehensive understanding of protein-antigen interactions^2^.

Surface plasmon resonance (SPR) and biolayer interferometry (BLI) are widely employed techniques that enable label-free, real-time analysis of binding kinetics^3,4^. These methods provide detailed kinetic profiles by measuring both association and dissociation phases of binding events.

However, conventional SPR and BLI assays require highly purified ligands and analytes, necessitating extensive preparation that increases time and resource requirements^3,5^. Additionally, these techniques demand meticulous control over ligand immobilisation on sensor surfaces. Insufficient ligand loading can lead to poor signal-to-noise ratios, while excessive loading causes surface heterogeneity, hindering accurate curve fitting and introducing artefacts such as mass-transport effects^6^. Disordered ligand immobilisation – caused by random ligand orientations following attachment to sensors – can further exacerbate these challenges^7,8^. This issue commonly arises when immobilisation to the sensor relies on methods such as protein adsorption, amine-mediated covalent attachment, or the use of streptavidin-coated sensors with ligands biotinylated at random amine groups, all of which typically results in a range of ligand orientations and hence different exposures of the binding sites.

The ability to accurately measure binding kinetics efficiently and cost-effectively, using minimal amounts of binding proteins such as nanobodies or single-chain variable fragments (scFvs) without purification steps, is highly desirable. This need is amplified by the rise of computational protein design and optimisation techniques^9–14^, as well as the establishment of next-generation sequencing (NGS) as standard to characterise panned libraries in antibody and binding-protein discovery^15–18^. These approaches yield large numbers of potential binders for the same target, necessitating high-throughput methods for characterisation. Rapid experimental feedback is crucial to enable iterative design cycles in computational approaches, and the ability to accurately screen numerous binders from NGS hits accelerates the identification of candidates with desired properties. In these contexts, obtaining a single antigen at high purity is entirely feasible. However, purifying each individual binder from these approaches is often impractical due to significant time and resource requirements, which typically constrains the number of binders that are characterised in the lab.

Efforts to utilise BLI or SPR with non-purified binders have been reported, with BLI being particularly favoured due to its disposable biosensors and higher throughput capabilities^19–22^. In these approaches, raw binding proteins or antibody fragments, such as nanobodies or scFvs, are typically contained in bacterial periplasmic extracts or mammalian cell supernatants. Two main strategies exist for kinetic measurements using such non-purified samples. The first involves immobilising the purified antigen on the sensor as the ligand and introducing the non-purified antibody fragments as the analyte in solution. However, unknown concentrations of binders in crude mixtures necessitate additional quantitation assays, increasing time and resource requirements. Without known concentrations, fitting the association rate becomes unfeasible, limiting analysis to the dissociation phase, whose rate is independent of analyte concentration, an approach that has been referred to as “off-rate screening”^7^. Moreover, non-specific binding from impurities in the analyte crude mixture can generate spurious signals, complicating data interpretation and curve fitting for both association and dissociation phases^6^.

The second strategy captures the antibody fragment from a crude mixture directly onto the sensor, using pre-coated sensors specific to purification tags (e.g., anti-His-tag sensors) or domains (e.g., anti-Fc-domain sensors). The purified antigen, at known concentrations, serves as the analyte. While this approach mitigates some issues related to unknown concentrations in crude mixtures, it introduces other limitations. Accurate measurement of the antibody-antigen interaction requires that the dissociation rate of the antigen (i.e., of the analyte from the ligand) is faster than the background dissociation rate of the captured antibody fragment from the sensor (i.e., of the loaded ligand) thus limiting the applicability range. Additionally, variable concentrations of different antibody fragments in crude extracts make it challenging to optimise ligand loading uniformly across multiple sensors. Overloading can lead to surface heterogeneity and mass transport artefacts, while underloading results in inadequate signal strength^6,22^. The necessity for consistent loading times across sensors when characterising multiple binders in parallel further complicates the assay, since differing concentrations in the samples can cause inconsistent results. Addressing this issue requires conducting preliminary quantitation experiments to standardise ligand concentrations, thus increasing both cost and time.

In this work, we introduce the SpyBLI method that overcomes all these limitations, enabling accurate quantification of binding kinetics from non-purified binders at unknown concentrations. Our approach eliminates the need for purification and concentration determination of ligands. Additionally, we demonstrate that a single BLI sensor can be employed to probe multiple analyte concentrations without excessively sacrificing accuracy, further reducing costs and enhancing throughput. This approach is usually referred to as single-cycle kinetics in SPR experiments, but it is not commonly implemented in BLI, possibly because the software for most BLI systems is set up for multi-cycle analysis. To overcome this limitation, we make available an easy-to-use Jupyter Notebook to process exported BLI raw data and perform single-cycle kinetics analysis with various fitting models.

We also leverage advances in cell-free expression to show that accurate binding kinetics can be obtained in less than 24 hours from receiving inexpensive linear gene fragments encoding the binders of interest, using as little as 10 µL of cell-free reaction mixture. The ability to obtain binding kinetic data rapidly and efficiently holds substantial promise for improving success rates and enhancing the chances of obtaining high-affinity binders ideally suited for downstream applications in research, diagnostics, or therapeutics.

## Results

### Binding kinetics quantification from non-purified binders

We introduce a new method to quantify binding kinetics that combines the synthesis of linear gene fragments, with cell-free expression systems or medium-throughput Golden Gate Cloning and mammalian expression^23^, the SpyTag003/SpyCatcher003 rapid covalent reaction^24^, and biolayer interferometry.

To initiate the process, gene fragments encoding the binders of interest are ordered from commercial suppliers as linear DNA fragments. In our study, we utilised two types of fragments (**Fig. 1A**). The first type contains sequences codon-optimised for mammalian expression, flanked by Golden Gate restriction sites. These fragments facilitate rapid one-step cloning into a mammalian expression vector that includes a CD33 secretion signal at the N-terminus and appends SpyTag003 and His-tag sequences at the C-terminus (see Methods). The second type comprises longer gene fragments forming the minimal gene expression unit for cell-free expression, incorporating a T7 promoter, a ribosome binding site, the binder sequence codon-optimised for bacterial expression, the SpyTag, and an optional His-tag for purification (**Fig. S1** and Methods). We find that these linear gene fragments can be directly introduced, without any cloning or PCR step, into *E. coli*-based cell-free expression systems to yield sufficient protein quantities for binding quantification.

**Figure 1.**
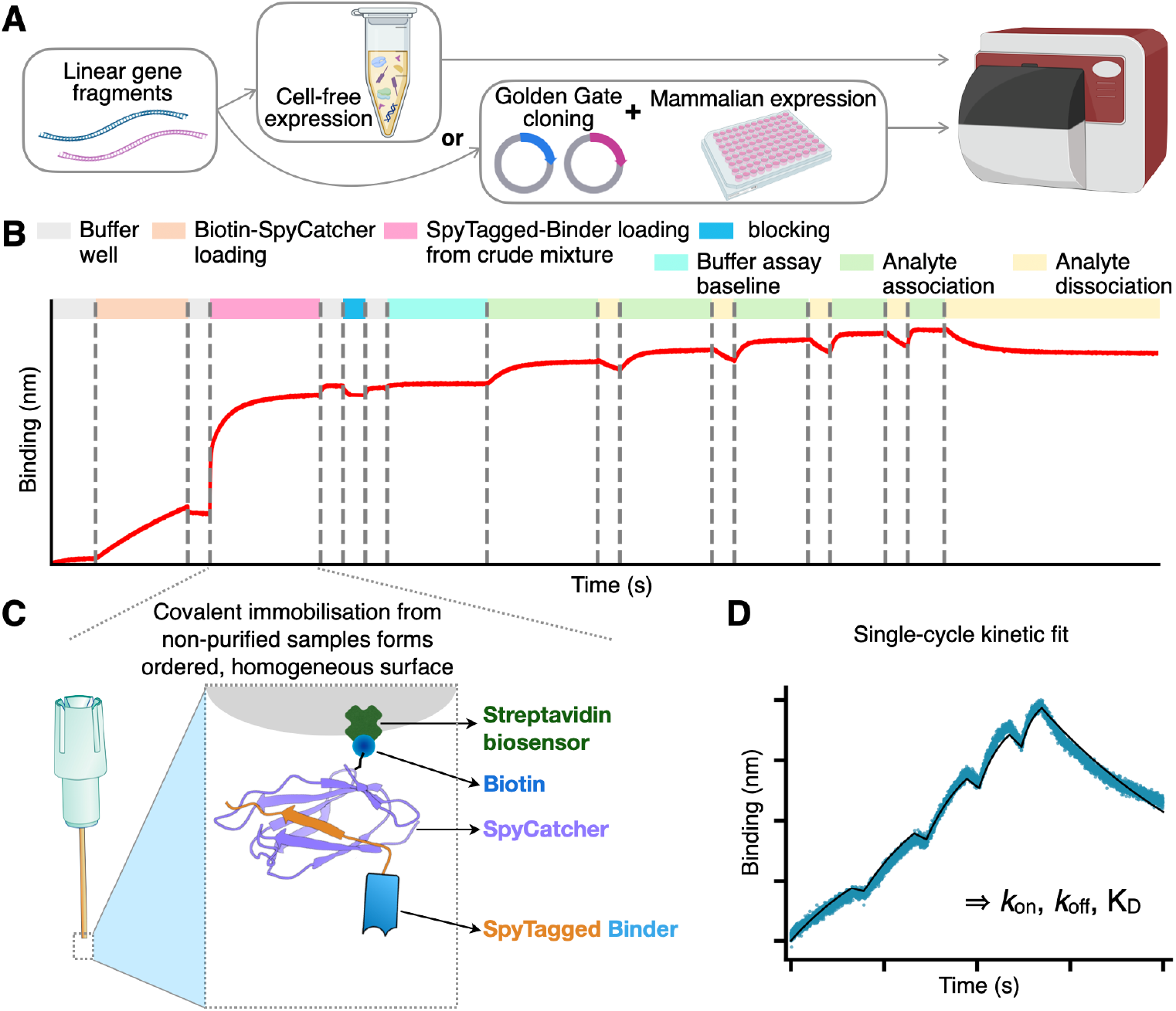
Overview of the SpyBLI pipeline. (**A**) Binders of interest are encoded in linear gene fragments, which are either used directly in cell-free expression, or Golden-Gate-cloned into vectors for expression in mammalian cell media. (**B**) Overview of the full BLI assay set up for a single assay sensor, with all steps highlighted (see legend) (**C**) Schematic of the fully loaded BLI assay sensor, forming a uniform surface of similarly oriented binders. (**D**) Example of a single-cycle kinetics binding curve (blue) obtained from an assay sensor probing increasing concentrations of antigen during the various association steps. This curve is fitted with a binding model (black line) to extract kinetic rate constants (*k*_on_, *k*_off_) and equilibrium dissociation constant (K_D_= *k*_off_ / *k*_on_).

Following expression, the binders – either present in crude mammalian cell supernatants or within cell-free expression mixtures – are directly utilised in a BLI assay (**Fig. 1B**). The assay employs streptavidin-coated sensors, onto which we load a predetermined amount of a purified S49C variant of SpyCatcher003, selectively biotinylated at the solvent-exposed engineered cysteine residue using maleimide chemistry (see Methods). This 1:1, site-specific biotinylation ensures a highly ordered sensor surface, with all SpyCatcher003 molecules predominantly oriented in the same manner. Using purified SpyCatcher003 at known concentrations allows precise control over the loading process, ensuring that all sensors possess a comparable density of SpyCatcher003 sites.

Subsequently, the sensors are immersed in wells containing the unpurified binders. The interaction between SpyCatcher003 and SpyTag003 is highly specific and rapid, enabling efficient covalent coupling through an isopeptide bond even when binder concentrations are low^24^. Importantly, the irreversible nature of this covalent interaction guarantees that once the binders are immobilised, they do not dissociate from the SpyCatcher003 molecules. Additionally, since all binders feature a C-terminal SpyTag003, the uniform orientation of ligands on the sensor surface is maintained (**Fig. 1C**). The sensors are loaded to saturation; wells with highly expressing binders achieve saturation swiftly, while those with lower expression levels take longer (**Fig. S2**). Nevertheless, due to the covalent bonding, given enough loading time all sensors will ultimately attain an equivalent density of immobilised binders, matching that of the pre-loaded SpyCatcher003 sites and ensuring uniformity across sensors. The signal observed during this binder-loading step also provides an opportunity to rank binders based on their expression levels, which is valuable information for binder characterisation and selection (**Fig. S2** and **S3B**,**C**).

After loading, the sensors are transferred to buffer wells to dissociate any non-specifically bound impurities. A brief blocking step follows, employing a high concentration of purified SpyTag003 peptide (**Supplementary dataset 1**). This blocking step has minimal impact on sensors already loaded with binders but is beneficial for control sensors used to monitor any non-specific binding of the analyte. The peptide effectively blocks and stabilises unoccupied SpyCatcher003 molecules, rendering the control sensors more comparable to the assay sensors, where the SpyCatcher003 is typically covalently bound to the SpyTag003 on the ligand.

After a brief wash, the sensors are transferred into the same kinetic buffer used for the analyte to establish a stable assay baseline. The robust biotin–streptavidin interaction^25^, coupled with the covalent SpyTag003–SpyCatcher003 bond, typically leads to a flat baseline (**Fig. 1B**), reducing the need for reference subtraction. We find that employing a reference sensor – loaded similarly but monitoring only buffer wells – is often unnecessary if the baseline is stable. Although using it may slightly refine kinetic parameter fits, we have not used reference subtraction across this study to maximise the number of sensors available for binder characterisation. However, we recommend including a blocked SpyCatcher003-loaded sensor once per antigen concentration series, to check for any non-specific binding of the analyte to the sensor.

The kinetic measurements proceed by transferring the sensors into antigen wells containing increasing concentrations of the analyte, typically prepared through serial dilutions (e.g., 1:2 or 1:3). Short dissociation steps are interspersed between association phases, culminating in a final, extended dissociation phase in buffer. The data collected are then fitted using an appropriate binding model to extract the kinetic parameters *k*_on_ and *k*_o,_, and equilibrium K_D_ (**Fig. 1D**).

To demonstrate the utility and reliability of our method, we applied it to a range of nanobodies and scFvs. We first selected the anti-β_2_-microglobulin nanobody Nb24^26^ and an anti-CD16a scFv, which corresponds to the FcγRIIIa-targeting arm of the bispecific antibody RO7297089^27^. **Figure 2** shows that the resulting binding sensorgrams are highly consistent whether using antibodies that have undergone extensive purification – consisting of affinity chromatography followed by size-exclusion chromatography – or antibodies obtained directly from crude mammalian cell supernatants or cell-free expression mixtures (**Fig. S4**). This consistency confirms that our method can reliably quantify binding kinetics without the need for binder purification. We further note that the K_D_ values we obtained (**Table 1**) are consistent with previously reported values of single-digit nanomolar for the anti-CD16a scFv and mid-nanomolar range for Nb24^26–28^.

**Table 1.** Binding kinetics parameters of the characterised nanobodies and scFvs. The table reports information on the various SpyTagged binders characterised in this study. The column ‘Source’ describes from where the binder was loaded on the sensor, either as a purified protein (Pure), or directly from mammalian cell supernatant (MCS) or from a cell-free reaction (CFR) blend. Literature K_D_ values are extracted from the given references, while measured parameters represent averages ± standard deviations over three independent experiments. Experiments for Iexkizumab and Secukinumab scFvs were carried out only once due to constraints in reagent and instrument availability. All sequences are provided in **Supplementary dataset 1**.

| Binder | Source | PDB ID | Antigen | Literature $K_D$<br>& Ref. | Measured $K_D$ | $k_{on} \times 10^5$<br>($M^{-1}s^{-1}$) | $k_{off}$<br>($s^{-1}$) |
| --- | --- | --- | --- | --- | --- | --- | --- |
| | Pure | | | | $49.6 \pm 8$ nM | $2.3 \pm 0.1$ | $(1.12 \pm 0.19) \times 10^{-2}$ |
| **Nb24** | MCS | 4kdt | $\beta_2m$ | $58$ nM <sup>36</sup> | $50 \pm 10.3$ nM | $2.4 \pm 0.6$ | $(1.14 \pm 0.13) \times 10^{-2}$ |
| | CFR | | | | $63.8 \pm 5$ nM | $2.20 \pm 0.22$ | $(1.4 \pm 0.2) \times 10^{-2}$ |
| **anti-CD16a**<br>**scFv** | Pure | | | | $6.7 \pm 1.3$ nM | $1.71 \pm 0.09$ | $(1.13 \pm 0.16) \times 10^{-3}$ |
| | MCS | 7seg | CD16a<br>(FCGR3A) | $7.7$ to $18.4$ nM <sup>27</sup> | $4.3 \pm 0.4$ nM | $1.87 \pm 0.29$ | $(0.81 \pm 0.19) \times 10^{-3}$ |
| | CFR | | | | $8.1 \pm 0.8$ nM | $2.2 \pm 1$ | $(1.8 \pm 1) \times 10^{-3}$ |
| **Nb.b201** | MCS | 5vnw | HSA | $431 \pm 12$ nM <sup>29</sup> | $772 \pm 163$ nM | $3.2 \pm 1.5$ | $0.24 \pm 0.08$ |
| **Ixekizumab**<br>**scFv** | MCS | | IL-17A | $\leq 3$ pM <sup>33</sup> | $\sim 3$ pM | 13 | $3.4 \times 10^{-6}$ |
| **Secukinumab**<br>**scFv** | MCS | | IL-17A | $60 \pm 20$ pM <sup>35</sup> | $\sim 53$ pM | 16 | $8.3 \times 10^{-5}$ |
| **Nb cAb-Lys3** | CFR | 1mel | Lysozyme | $5 \pm 4$ nM <sup>37</sup> | $0.67 \pm 0.06$ nM | $17 \pm 3$ | $(1.16 \pm 0.26) \times 10^{-3}$ |
| **HyHEL10**<br>**scFv** | CFR | 2znw | Lysozyme | $130$ pM <sup>31</sup> | $143 \pm 36$ pM | $23 \pm 14$ | $(3.59 \pm 0.29) \times 10^{-4}$ |

**Figure 2.**
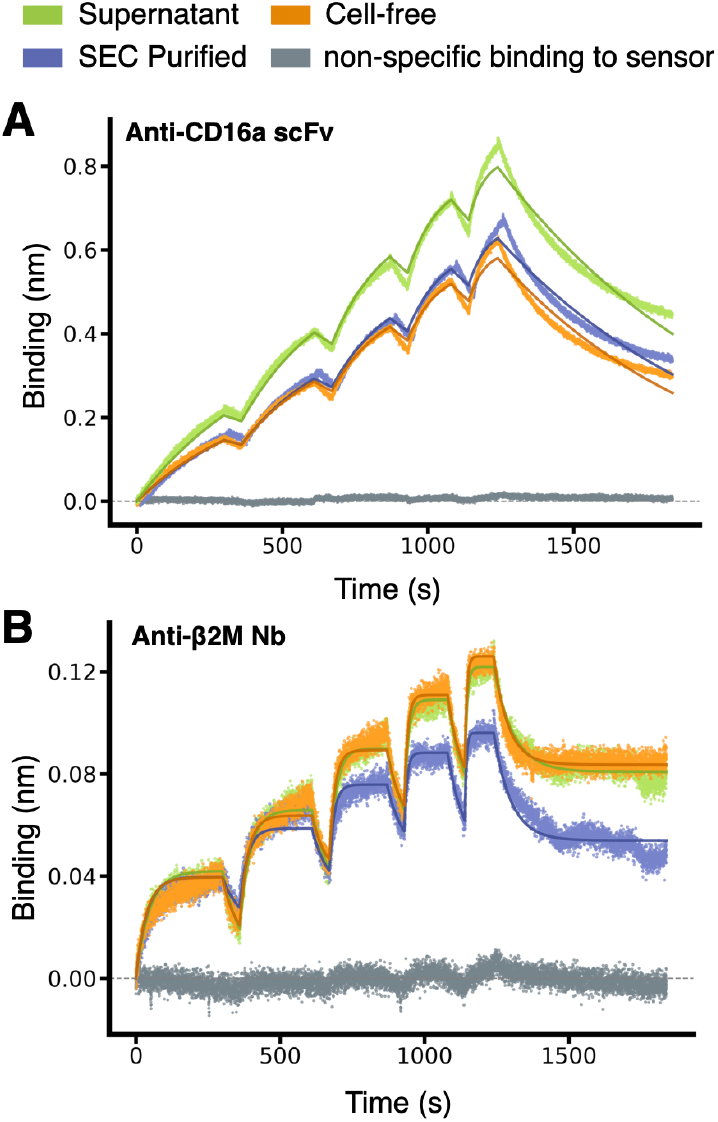
Consistency of binding kinetic measurements between purified and non-purified antibody fragments. (**A**) BLI sensorgrams of a scFv (PDB ID 7seg) binding to CD16a. The monovalent antigen was purified and used as analyte at increasing concentrations (6.25, 12.5, 25, 50, 100 nM) in each association phase. The SpyTagged scFv is used as ligand, and it was loaded either as purified scFv in buffer or from unpurified mixtures (see legend and **Fig. S4**). Different experiments (coloured lines) were carried out on different days, and the minor differences in R_max_ (maximum signal) observed can be rationalised by minor differences in loading. The solid lines correspond to a fit with a 1:1 standard binding model. (**B**) Same as **A** but for the nanobody Nb24 (PDB ID 4kdt) binding to purified β2-microglobulin used as analyte, which was present at 25, 50, 100, 200, 400 nM in each association phase, respectively. The solid lines correspond to a fit with a 1:1 partial dissociation binding model. Results of all fits are in **Table 1**.

Encouraged by these results, we extended our method to test additional nanobodies and scFvs across a broader range of binding affinities. We evaluated two more nanobodies: Nb.B201, which binds weakly to human serum albumin (HSA)^29^, and cAb-Lys3, which binds strongly to hen egg-white lysozyme^25^. Nb.B201 was expressed in mammalian cell supernatant, while cAb-Lys3 was produced using the cell-free expression system. As anticipated, Nb.B201 exhibited rather weak binding to HSA, with a K_D_ in the high nanomolar range, in agreement with literature values^29,30^ (**Fig. 3** and **Table 1**). Conversely, cAb-Lys3 demonstrated strong binding to lysozyme, with a K_D_ in the high picomolar range, consistent with previous reports (**Fig. 3** and **Table 1**).

**Figure 3.**
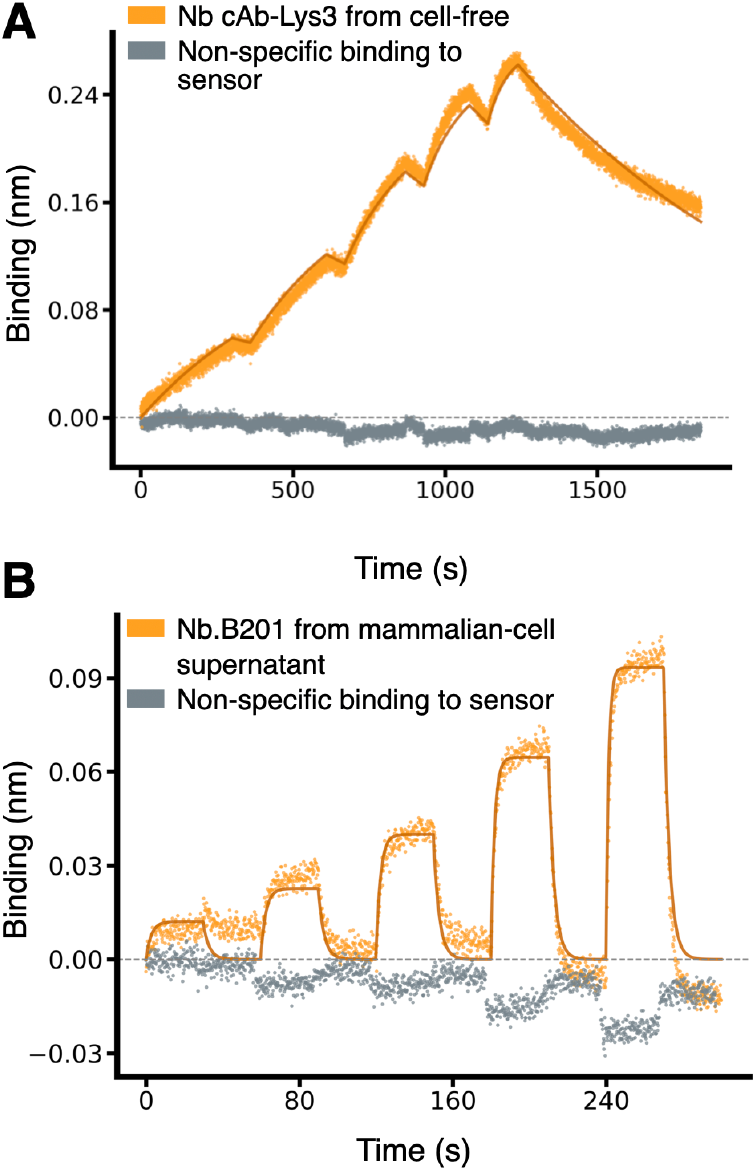
Characterisation of nanobodies spanning a broad range of affinities. (**A**) Binding sensorgrams of SpyTagged nanobody cAb-Lys3 (PDB ID 1mel) loaded from cell-free-expression blend and binding to purified hen egg-white lysozyme, which was used as analyte at increasing concentrations of 0.625,1.25, 2.5, 5, 10 nM. The solid lines correspond to a fit with a 1:1 standard binding model. (**B**) Binding sensorgram of SpyTagged nanobody Nb.B201 (PDB ID 5vnw) loaded from a mammalian-cell supernatant and binding to HSA, which was used as analyte at increasing concentrations of 62.5, 125, 250, 500, 1000 nM. The solid lines correspond to a fit with a 1:1 standard binding model.

We further tested three scFvs with literature-reported K_D_ values spanning from high to low picomolar ranges. These included the mouse scFv HyHEL10, targeting hen egg-white lysozyme^31^, and two therapeutic antibodies approved for clinical use, which we expressed as scFvs: Secukinumab^32^ and Ixekizumab^33^, both targeting human interleukin-17A (IL-17A). The BLI sensorgrams obtained using non-purified material were fitted to yield K_D_ values within the expected ranges^33–35^ (**Table 1**), demonstrating our method’s capability to accurately quantify high-affinity interactions (**Fig. 4**).

**Figure 4.**
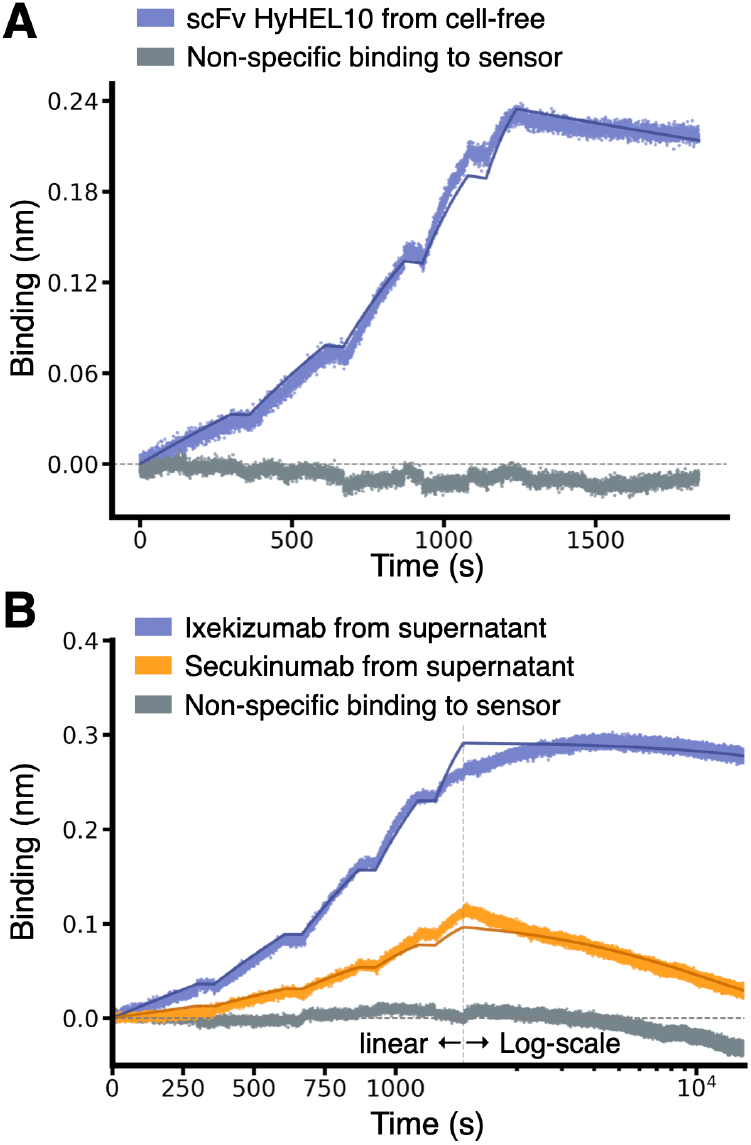
Characterisation of scFv binding in the pico-molar range. (**A**) Binding sensorgrams of SpyTagged scFv HyHEL10 (PDB ID 2znw) loaded from cell-free-expression mixture and binding to purified hen egg-white lysozyme, which was used as analyte at increasing concentrations of 0.31, 0.625, 1.25, 2.5, and 5 nM. The solid lines correspond to a fit with a 1:1 standard binding model. (**B**) Binding sensorgrams of SpyTagged scFv Ixekizumab and Secukinumab (see legend) loaded from a mammalian-cell supernatant and binding to IL-17A, which was used as analyte at increasing concentrations of 0.25, 0.5, 1, 2, and 4 nM. The association phase is plotted on a linear scale (x-axis), while the much longer dissociation phase on a log10 scale. The solid lines correspond to a fit with a 1:1 standard binding model.

We note that the single-digit picomolar affinity of Ixekizumab lies outside of the dynamic range reported for BLI (approximately 10 pM – 1 mM)^4,5^. In our data, the initial segment of the dissociation step appears to drift upward, and additional artefacts – potentially influenced by evaporation – emerge toward the end of the dissociation step, which had to exceed four hours in duration to see any dissociation (**Fig. 4B**). These artefacts underscore the inherent limitations of BLI for extremely tight binders. However, the fact that we could still fit a K_D_ value in the range of those obtained using SPR with purified proteins^33,34^ confirms that our covalent immobilisation strategy of non-purified binders does not restrict the technique’s intrinsic range.

Taken together, our results confirm that SpyBLI reliably quantifies binding kinetics across a wide spectrum of affinities, from high nanomolar to low picomolar K_D_ values, using unpurified binders that can be expressed in different systems. These findings underscore the versatility and robustness of SpyBLI, enabling rapid and cost-effective characterisation of diverse antibody fragments and, most likely, binding proteins more generally.

### Establishment of a small-scale cell-free expression system for gene fragments

For some of the antibody fragments examined, we relied on transient mammalian expression because we had them available in mammalian vectors. However, the setup for cell-free expression we have introduced offers unique advantages. By enabling the direct use of linear gene fragments, our approach eliminates the need for cloning or any PCR assembly step. Linear gene fragments can be added directly to the expression blend, enabling the measurement of binding kinetics immediately after overnight expression. In contrast, mammalian expression requires up to 10 days for cloning, transfection, and sufficient expression. This rapid turnaround time makes cell-free expression particularly appealing for early-stage screening and optimisation workflows.

To establish a robust *E. coli*–based cell-free expression system for nanobodies and single-chain variable fragments (scFvs), we optimised both the composition of the cell-free blend and the design of the linear DNA fragments. The reducing cytosolic environment of *E. coli* typically hinders the formation of disulfide bonds, which are vital for the correct folding and function of these binders – especially as two of the nanobodies we tested contain non-canonical disulfide bonds that stabilise the CDR3 conformation^38^. To address this challenge, we systematically explored combinations of additives and solubility tags using an automated eProtein Discovery instrument (see Methods), where we conducted two experiments. In the first, we screened a panel of four nanobodies with non-canonical disulfide bonds and two scFv variants. Each was tested with three different solubility tags as well as without a solubility tag, resulting icn a total of 24 DNA fragments. We then selected 8 different cell-free blends, each customized with the addition of two additives. For the first experiment, the selected additives included a mix of TrxB1 (thioredoxin reductase), a DnaK mix (a combination of molecular chaperones), a cofactor mixture, GSSG (oxidized glutathione), and PDI (protein disulfide isomerase). The results showed that omitting solubility tags and increasing the oxidative power of the cell-free blends improved production levels (**Fig. S5A**). We then performed a second experiment where we tested six additional nanobodies without solubility tags and expanded the assessment of the best oxidative conditions. The results show that increasing the oxidative power even further with the addition of high concentrations of PDI and GSG, further increases soluble yields of all the proteins tested (**Fig. S5B**). We then scaled up the expression, purified the 12 antibody fragments, and confirmed by mass spectrometry the correct formation of the disulfide bonds for all antibody fragments (**Table S1**).

In these experiments, the linear DNA fragments preparation required a one-step PCR, a DNA purification step, and a concentration-normalization step. We therefore looked at reducing the DNA preparation steps by creating a bespoke linear DNA construct that – after resuspension – could be used directly in the cell-free blend for the SpyBLI workflow. Eight different linear DNA fragments encoding SpyTagged Nb24 were designed, each differing in their 5′ and 3′ flanking regions, allowing us to test two 5’ end lengths, two post-promoter spacer lengths (between the T7 promoter and start codon), and the presence or absence of a T7 terminator, while keeping the coding sequence identical. All eight constructs yielded protein, albeit to varying extents. The most influential factor on expression level was the length of the region between the T7 promoter and the start codon, whereas alterations to the other regions had minimal impact (**Fig. S3**). Subsequent BLI analysis confirmed that each Nb24-SpyTag003 construct maintained expected binding kinetics and affinities for β2-microglobulin (**Fig. S3D**). The best-performing construct featured a short 5′-end region, followed by the T7 promoter, then a 47-nucleotide spacer, which we took from the pDEST *E. coli* expression vector used to express the SpyCatcher003 protein, and that contains the ribosome binding site, followed by the coding sequence and a short 3′-end region of just 10 nucleotides (**Fig. S1**).

Taken together, the results presented in this study demonstrate that our optimised cell-free system can reliably and rapidly produce functional nanobodies and scFvs, including those requiring non-standard disulfide bonds to stabilise the binding surface, such as Nb24 and cAb-Lys3. Cell-free-expressed antibody fragments yielded binding kinetics fully comparable to those from antibody fragments expressed in mammalian cells.

## Discussion

By leveraging the strengths of the biotin-streptavidin and SpyTag003/SpyCatcher003 interactions, we have presented a BLI-based method, called SpyBLI, that provides reliable and accurate kinetic measurements without requiring binder purification or concentration determination. We confirmed the method’s applicability across six orders of magnitude in affinity values, obtaining results that align well with previously reported data.

The uniform loading and orientation of binders on the sensor surface, along with the near elimination of ligand dissociation yield high-quality kinetic data. In principle, SpyCatcher003 S49C could be covalently conjugated directly on the sensor, thereby removing the need for an additional streptavidin layer that might contribute to non-specific binding. However, while streptavidin biosensors are commercially available, both in their standard form (used in this study) and various high accuracy formats, there are no commercially available biosensors for thiol conjugation, while amine conjugation would result in a disordered ligand orientation. Consequently, our current setup offers the best compromise among off-the-shelf availability, ease of use, and consistent binder orientation.

In this work, we have employed single-cycle kinetics, in which a single BLI sensor probes multiple analyte concentrations, since this approach enables higher throughput and reduces sensor usage and hence costs. However, our method is fully compatible with more traditional multi-cycle-kinetics BLI protocols, in which different sensors are employed to probe different analyte concentrations, and their signal is then fitted globally to determine the binding rate constants. We have not explored the method’s applicability to mini-proteins^11,13^ and other antibody mimetics^39–43^, but we would expect these to be easier than antibody fragments, as they are typically highly stable, easy to express, and devoid of disulfide bonds.

We further streamlined our approach by integrating advances in cell-free expression and by optimising the design of linear gene fragments, which allowed us to determine binding kinetics in less than 24 hours of receiving inexpensive gene fragments. This protocol requires minimal hands-on time and has no cloning or PCR steps. We showed that both scFvs and nanobodies – including those containing non-canonical disulfide bonds – produced in cell-free expression blends display binding constants consistent with those obtained from crude mammalian cell supernatants or purified with affinity and size-exclusion chromatography.

One potential limitation of our cell-free approach is that linear gene fragments are not entirely error-free. Although the typical error rate for this type of DNA synthesis is below one in 5,000 base pairs^44–46^, our scFv-expressing fragments have approximately 950 bases, about 750 of which are the protein-coding sequence. Therefore, one may expect at least one error in up to 14% of the fragment pool encoding an scFv. In fragment synthesis the most common errors are single-nucleotide deletions^44,46^, which, like insertions, would disrupt the reading frame and prevent the correct translation of the SpyTag003 at the C-terminus, thereby preventing any frame-shifted product from loading on the sensor. In contrast, single-nucleotide substitutions, when non-synonymous, may produce proteins that still load onto sensors but contain mutated amino acids. Nevertheless, this mutated population should represent a small minority (under 10% for scFvs and even less for nanobodies), and only a fraction of possible mutations would affect binding affinity. While the presence of such sub-population may introduce some heterogeneity into the binding traces, we have not observed any adverse effects on the reliability of our kinetic measurements. Cell-free expression data remain consistent with those obtained from sequence-verified mammalian expression, indicating that any potential distortion from error-containing fragments is undetectable within the noise intrinsic in BLI measurements.

In conclusion, the SpyBLI approach we have introduced provides a rapid, cost-effective, and high-throughput solution for accurately quantifying binding kinetics directly from unpurified samples, thereby accelerating the characterisation of candidate binders. We anticipate that SpyBLI will be especially valuable in computational protein design^9–14^, binder optimisation^47–50^, and the high-throughput screening of binding candidates identified through next-generation sequencing of panned libraries^15–18^.

## Materials and methods

### Gene synthesis and antibody fragment mammalian expression

DNA sequences encoding the selected nanobodies and scFvs were ordered as gene fragments (Gene Titan platform, GenScript), either with human-optimised codons and containing Golden Gate cloning sites for insertion into a mammalian expression vector, or with *E. coli*-optimised codons as full linear expression fragment for cell-free expression (see later). Codon optimisation was performed using the online optimization tool from GenScript. Amino acid sequences were retrieved from the Protein Data Bank (see PDB ID in captions), except those of the therapeutic antibodies that were retrieved from Thera-SAbDab^51^. All sequences can be found in **Supplementary dataset 1**.

For mammalian expression, gene fragments were cloned using Golden Gate BsmBI-v2 kit (New England Biolabs; E1602S) in a pcDNA3.4 mammalian expression vector. The vector was modified to contain an N-terminal CD33 secretion sequence and a C-terminal SpyTag003 followed by a 6xHis tag. Furthermore, gene fragments were designed to have a (G_3_S)_2_ linker between the nanobody or scFv domain and the SpyTag003, to reduce any steric hindrance in the interaction with SpyCatcher003.

Cloned plasmids were transformed into DH5*α* competent cells (New England Biolabs, #C2987H) and grown overnight at 37 °C on LB media plates containing ampicillin before midi prep cultures were set up the next day. Midi preps were processed using QIAGEN Midi Prep kit (QIAGEN). Purified plasmids were sent for Sanger sequencing and, upon confirmation of the correct sequence, were used for protein expression.

Plasmids were transfected into Expi293F cell line following protein transfection protocol from the manufacturer (ThermoFisher Scientific; A14635). For nanobody and scFv expression 3 mL cultures were set up. Cells were incubated for 3 days at 37 °C with 5% CO_2_ on an orbital shaker with 120 rpm. On day 3, cells were harvested by centrifugation (4 °C, ∼2700 g, 20 minutes) and the supernatant was either used for protein purification or directly in the BLI assays.

### Antigen preparation

CD16a-mMBP sequence was designed as described in Ref. ^52^, codon-optimised for mammalian expression, and ordered as a gene fragment from Twist Bioscience. It was cloned into pcDNA3.4 vector that did not contain SpyTag003 sequence using BsmBI-v2 Golden Gate assembly kit (New England Biolabs; E1602S). Resulting plasmids were confirmed by sequencing and transfected into 30 mL cultures (Expi293F cell line). Cultures were harvested 6 days post-transfection as described above.

Recombinant β2-microglobulin was expressed and purified to homogeneity as reported in Ref. ^53^. Human Serum Albumin was purchased from Sigma-Aldrich (A3782) as lyophilized powder. It was reconstituted in PBS and purified by size exclusion chromatography (SEC) using a Superdex 200 Increase 10/300 GL column, prior to being used in BLI assays against Nb.B201. Lysozyme from chicken egg-white (Sigma-Aldrich; 62971) was reconstituted in PBS and purified by SEC using a Superdex 75 Increase 10/300 GL column.

### Protein purification

His Mag Sepharose Excel magnetic beads (Cytiva) were washed with PBS before being added to mammalian-cell supernatants. For each culture, 0.1 mL to 0.5 mL of settled beads was added, and samples were incubated on a roller at 4 °C for 2-3 hours. Beads were washed and resuspended in PBS to be processed on AmMag™ SA Plus Semi-automated System 980 (Genscript). In the system, beads are washed with PBS and 4mM Imidazole and eluted with 200mM Imidazole. Eluted proteins are further purified by SEC on an AKTA Pure system to remove the Imidazole and isolate the monomeric protein. A Superdex 75 increase 10/300 GL column was employed for proteins with MW < 50 kDa and a Superdex 200 increase 10/300 GL column for the others. PBS was used as a running buffer. Resulting purified proteins in PBS were aliquoted and flash frozen in liquid nitrogen. Samples were stored at -80 °C.

### SpyCatcher003 S49C expression and purification

SpyCatcher003 S49C was obtained in pDEST114 plasmid^24^ (Addgene #133447) and transformed into *E. coli* C41(DE3) cells (Merck; CMC0017). Colonies were grown on an ampicillin agar plate at 37 °C overnight. A colony was picked to set up an overnight 10 mL culture in a shaking incubator (180rpm, 37 °C). Next day, some of the sample was taken to make a glycerol stock, with the rest being added to 1L flask of LB media supplemented with 100 µg/mL ampicillin and returned to a shaking incubator to allow cells to grow. When optical density (OD) reached 0.6, IPTG was added at a final concentration of 0.42mM and the flask was incubated overnight at 28 °C (200 rpm). Next day, the culture was spun down at 6,000 g for 20 minutes. Supernatant was discarded and cell pellet was resuspended in lysis buffer (PBS + EDTA-free protease inhibitor tablet). Resuspended pellet was sonicated on ice (15 s on/45 s off; 20 minutes total). After sonication, lysed cells were centrifuged at 20,000 g for 30 minutes. Resulting supernatant containing SpyCatcher003 S49C was filtered with 0.45 µm PES membrane filter (Merck Millipore; SLHP033RS). Then, His Mag Sepharose Excel magnetic beads were added to perform IMAC purification, followed by SEC following the steps described above in ‘protein purification’.

Post-SEC, the His tag was removed by cleavage with TEV protease (New England Biolabs; #P8112S) following manufacturer instructions. The cleavage reaction was carried out at RT for 4-5 hours on a roller. After cleavage, the sample was incubated for 1 hour with His Mag Sepharose beads to remove cleaved His tags and any uncleaved SpyCatcher003 S49C. Beads were then removed by centrifugation and the resulting supernatant was size excluded again. Successful cleavage was confirmed by liquid-chromatography mass spectroscopy using VION (Waters, **Fig. S6**). We note that His tag cleavage is not strictly necessary to run the SpyBLI pipeline. However, we also use this reagent for other assays that would be hindered by the presence of a His tag onto the capturing SpyCatcher003 molecule. Therefore, the S49C SpyCatcher003 used in this work always had the His tag removed – a procedure that also further increased purity because of the additional purification steps.

#### SpyCatcher003 S49C biotinylation

TEV-cleaved SpyCatcher003 was biotinylated at the engineered cysteine site at S49C using EZ-Link Maleimide-PEG2-Biotin (Thermo-Fisher; A39261) following manufacturer instructions. The reaction was carried out at 4 °C overnight on a roller, and after centrifugation at 4 °C for 10 minutes at maximum speed on a benchtop centrifuge to pellet down any precipitate. SEC was then used to remove free biotin and to further purify the protein, as disulfided dimers may form during the labelling reaction. Complete 1:1 biotinylation was confirmed by liquid-chromatography mass spectroscopy using VION (Waters, **Fig. S6**).

#### Nuclera eProtein Discovery system

To setup the cell-free expression of SpyTagged scFvs and nanobodies, we first optimised the reaction conditions using the eProtein Discovery system. We performed cell-free expression of various nanobodies and scFvs to refine the cell-free blends components and determine the most suitable solubility tags, if any. First, DNA coding sequences of interest were designed and codon optimized directly in the eProtein Discovery software. The sequences included two small flanking sequences encoding for 3C and TEV proteases cleavage sites used in the subsequent overlapping PCR reactions. The sequences were ordered as gBlocks™ from Integrated DNA Technologies. One-step overlapping PCRs were carried out to assemble linear expression cassettes from ordered gene fragments. In this way, regulatory elements like promoter and terminator, solubility tag, detection tag and Strep tag were added to the coding sequence following the eGeneTM Prep Kit User Guide for the Solubility Tag Screen kit (Nuclera, NC3009). The one-step PCR was assembled adding the gBlock, the provided left megaprimer (containing promoter, RBS, translation enhancer, solubility tag), the provided right megaprimer (detector tag, Strep-tag, terminator). The assembled eGenes were purified and normalized to 5nM concentration with the eGene elution buffer and used to run the eProtein Discovery screen (Cartridge Reagent kit NC3010). Each eGene was expressed with different cell-free blends to identify the optimal expression conditions for our proteins. The screenings were set up following the instrument’s step-by-step guide.

From this screening, we identified that the highest-yielding constructs were those without any solubility tag and expressed with the addition of the GSSG/PDI additive (**Fig. S5)**. The lowest binder concentration observed in the cell-free blend was approximately 13 µM. Assuming similar expected yields for other linear gene fragments, we concluded that adding only 2 µL of the cell-free blend post-expression to a final volume of 200 µL, which is the volume required in a BLI assay well, would achieve at least 100 nM SpyTagged protein concentration. According to the results in **Fig. S2**, this concentration is sufficient for loading onto the sensor in a reasonable timescale. Therefore, we followed this dilution strategy when using cell-free-expressed binders in the SpyBLI assay.

### Cell free protein expression from gene fragments

Linear DNA fragments encoding for Spy Tagged nanobodies or scFvs of interest were designed and ordered from GenScript (Gene Titans). These fragments included a 5’-end T7 promoter, spacer and Ribosome Binding Site (see **Fig. S1**). Cell-free protein expression reactions were set up for overnight incubation (17 hours) at 29 °C using Nuclera Scale-Up kit following corresponding protocols for cell-free reactions from Nuclera (NC3004, NC3005). Each reaction was set up in a total volume of 20 µL. Expression of the desired proteins was confirmed the next day by running samples on Sodium dodecyl sulphate polyacrylamide gel electrophoresis (SDS-PAGE) stained with InstantBlue™ Protein Stain (Sigma-Aldrich) and by the observed loading traces on the BLI.

#### Biolayer Interferometry

All assays were performed on an Octet-K2 BLI system (Sartorius), except for those involving Secukinumab and Ixekizumab scFvs, which were performed on an Octet-Red BLI system (ForteBio), and the experiment in **Figure S7**, which was conducted on a GatorBio BLI system (GatorBio). All runs were performed at 30 °C with 1000 rpm shaking in PBS pH 7.5 with 0.05% (v/v) Tween-20. Assays were set up in 96-well plates (Greiner 655209) with 200 µL per well. Streptavidin biosensors (Sartorius 18-5019) were pre-hydrated in the running buffer for at least 15-20 minutes before the run.

Biotinylated S49C SpyCatcher003 was loaded at a concentration of 12.5 nM, except for the assay in **Figure S7**, which used varying concentrations. To ensure optimal binding kinetics and avoid overloading the sensor, the loading time of biotinylated S49C SpyCatcher003 was typically adjusted to load a total response of maximum 0.15 nm. **Figure S7** presents a dedicated experiment to systematically assess the effect of different SpyCatcher003 loading amounts. As expected, the results show that the signal to noise increases the more SpyCatcher003 is loaded, but so do deviations from the theoretical binding models, which likely result from surface heterogeneity and mass-transport artifacts that become more pronounced with increased crowding of the sensor surface.

In assays using purified SpyTagged003 proteins, a loading concentration of 100 nM was used (except for the assay in **Figure S2**, where this was systematically varied over more than 10 folds). For proteins loaded directly from mammalian-cell supernatant, the supernatant was mixed at a 1:1 ratio with PBS + 0.05% (v/v) Tween-20. For proteins expressed in the cell-free system, the cell-free blends were diluted 100-fold with PBS + 0.05% (v/v) Tween-20 (2 µL of cell-free blend in 200 µL final volume).

Association phases were performed at increasing concentrations of the relevant antigen (see figure captions) with times of 300s, 250s, 200s, 150s, and 100s respectively for lowest to highest antigen concentration.

## Acknowledgments

We are grateful to Dr Katherine Stott for granting us access to a Gator BLI instrument as it was demonstrated in the Biophysics Facility at the Department of Biochemistry. Some panels in Figure 1 were created with the help of BioRender.com. P.S. is a Royal Society University Research Fellow (grant no. URF\R1\201461). We acknowledge funding from UK Research and Innovation (UKRI) Engineering and Physical Sciences Research Council (grant no. EP/X024733/1, an ERC starting grant to P.S. underwritten by UKRI). S.R. acknowledges funding from the AIRC foundation (grant no. IG 2024 ID 30307) and from Cariplo/Telethon Foundations (GJC23044). M.R.H. acknowledges funding from the Medical Research Council (MR/Y011910/1).

## Data Availability

All data needed to evaluate the conclusions in this article, or that are necessary to interpret, verify and extend the research in the article are present in the paper and/or the Supplementary Materials and Supplementary files. Additional details are available from the corresponding author on request. A Python Jupyter Notebook to pre-process exported raw BLI data, and carry out global fits of single-cycle kinetics obtained with a single sensor (as carried out in this work) is made available at: https://gitlab.developers.cam.ac.uk/ch/sormanni/bli_one_sensor_multi_concentration

## Author contributions

O.P., M. Atkinson and P.S. designed experiments. O.P., M. Atkinson, and M.Ali performed experiments. A.H.K. and M.R.H. advised on experiment design, provided reagents, expert advice and feedback. C.V. and S.R. produced β2-microglobulin and provided expert advice and feedback. P.S. conceived and supervised the project. O.P., O.W. and P.S. analysed the data and wrote the first draft of the paper. All authors edited the paper.

