## Supplementary Information for "The SpyBLI cell-free pipeline for the rapid quantification of binding kinetics from crude samples"

| Antibody fragment | Theoretical MW (Da) | Measured MW (Da) | Δ MW (Da) | Expected disulfide bonds |
|---|---|---|---|---|
| scFvH wt | 33977.37 | 33973 | 4.37 | 2 |
| scFvH opt | 34093.67 | 34090 | 3.67 | 2 |
| VHH OAS 843 6Cys | 22125.42 | 22119 | 6.42 | 3 |
| VHH sdAb 2505 Cd\|Arabian 4Cys | 21651.83 | 21648 | 3.83 | 2 |
| VHH sdAb 1510 Cb\|Bactrian 4Cys | 21372.46 | 21368 | 4.46 | 2 |
| sdAb 4875 Sy\|Synthetic 4Cys intra-domain | 21476.73 | 21473 | 3.73 | 2 |
| 5vxm B vicugna 4Cys | 21411.75 | 21407 | 4.75 | 2 |
| sdAb 8797 Ca\|Camelid 2Cys | 20893.15 | 20891 | 2.15 | 1 |
| 6itc V lama 2Cys | 20864.29 | 20863 | 1.29 | 1 |
| 6xxp A camelus 2Cys | 21375.43 | 21373 | 2.43 | 1 |
| 7kgj B synthetic 2Cys | 21428.71 | 21426 | 2.71 | 1 |
| 3r0m B lama 2Cys | 21729.92 | 21728 | 1.92 | 1 |

**Table S1. Theoretical and measured molecular weights (MWs) of nanobodies and scFvs in cell-free blends.** Theoretical MWs were calculated from the amino acid sequences using the ProtParams tool (https://web.expasy.org/protparam/), which assumes reduced disulfides, whereas measured values were obtained via liquid chromatography–mass spectrometry with electrospray ionisation (see Methods). The antibody fragments are the same as those in **Figure S5** without solubility tags, and their sequences are provided in **Supplementary Dataset 1**. Prior to measurement, each protein was purified by strep-tag affinity chromatography. The "Δ MW" column indicates the difference between the theoretical and measured MW. Each correctly formed disulfide bond is expected to account for a 2 Da difference, and the expected number of disulfide bonds is listed next to it.

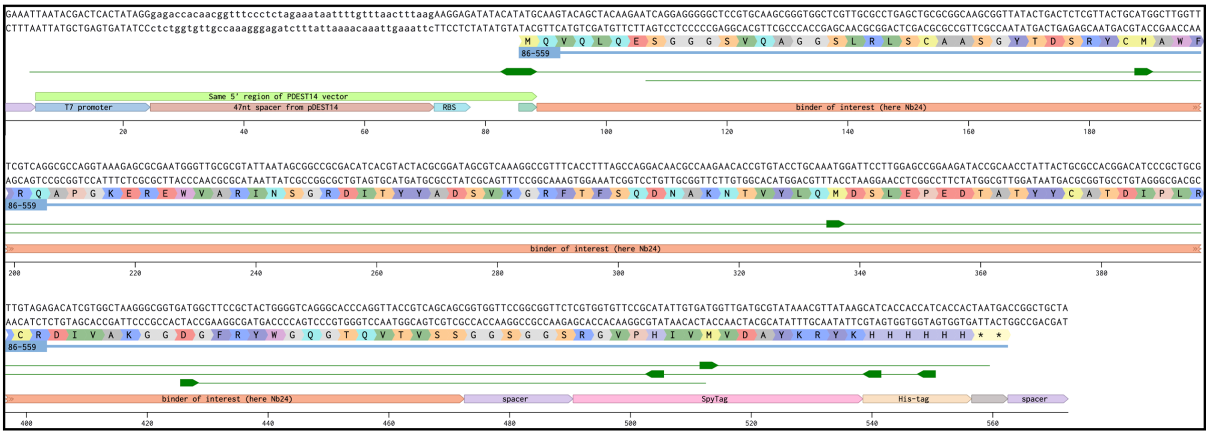


**Figure S1. Sequence of linear DNA fragment used for cell-free expression.** DNA sequence of the double-stranded linear DNA fragment optimised for cell-free expression. The coding sequence is also shown as translation. This example is for the Nb24 nanobody, but all other binders were obtained in the same way by substituting the binder sequence (orange annotation) with that of the binder of interest. The various annotations below the sequence describe what the relevant pieces are: RBS stands for ribosome binding site, and the His-tag at the C-terminus of the coding sequence is optional (it is not needed to run the SpyBLI assay, but it was present in the constructs used in this study, as it can be used for affinity-purification). We also provide a copy-paste friendly version of the DNA sequence, with the parts encoding Nb24 and the 6xHis-tag in lower case, and start and stop codons in bold font:

GAAATTAATACGACTCACTATAGGGAGACCACAACGGTTTCCCTCTAGAAATAATTTTGTTTAA
CTTTAAGAAGGAGATATACAT**ATG**caagtacagctacaagaatcaggagggggctccgtgcaagcgggtggct
cgttgcgcctgagctgcgcggcaagcggttatactgactctcgttactgcatggcttggtttcgtcaggcgccaggtaaag
agcgcgaatgggttgcgcgtattaatagcggccgcgacatcacgtactacgcggatagcgtcaaaggccgtttcacctt t
agccaggacaacgccaagaacaccgtgtacctgcaaatggattccttggagccggaagataccgcaacctattactgc
gccacggacatcccgctgcgttgtagagacatcgtggctaagggcggtgatggcttccgctactggggtcagggcaccc
aggttaccgtcagcagcGGTGGTTCCGGCGGTTCTCGTGGTGTTCCGCATATTGTGATGGTTGATG
CGTATAAACGTTATAAGcatcaccaccatcaccac**TAATGA**CCGGCTGCTA

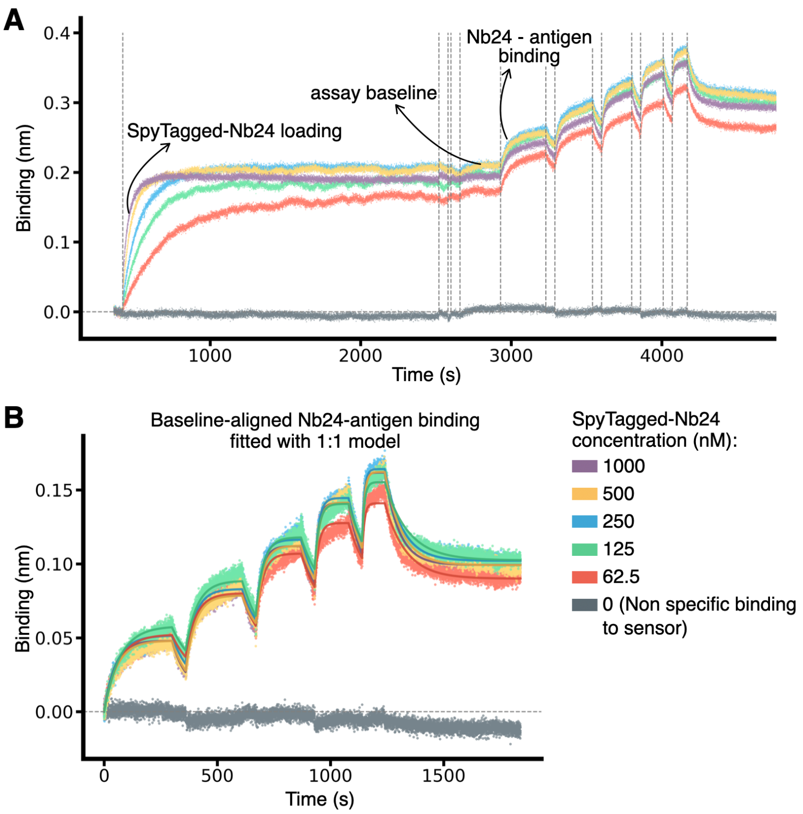


**Figure S2. Robustness of kinetic measurements across a more than 10-fold variation in loading concentration**. (**A**) BLI sensorgrams resulting from the loading of different concentration of SEC-purified SpyTagged Nb24 (see legend) onto sensors pre-loaded with biotinylated SpyCatcher003 and then probing the same wells as detailed in **Fig. 1** of the main text. (**B**) Fits carried out with a 1:1 partial dissociation binding model (solid lines) of the analyte ($\beta_2$microglobulin) associations and dissociations steps. Traces are aligned to the assay baseline, and the antigen concentrations were 25, 50, 100, 200, and 400 nM.

**A**

| Fragment ID | Length of 5' flanking site | Length of T7 promoter to ATG | T7 terminator |
|---|---|---|---|
| Nb24_fr.1 | 5 bp | 11 bp | yes |
| Nb24_fr.2 | 20 bp | 11 bp | yes |
| Nb24_fr.3 | 5 bp | 47 bp | no |
| Nb24_fr.4 | 20 bp | 47 bp | no |
| Nb24_fr.5 | 5 bp | 47 bp | yes |
| Nb24_fr.6 | 20 bp | 47 bp | yes |
| Nb24_fr.7 | 5 bp | 11 bp | no |
| Nb24_fr.8 | 20 bp | 11 bp | no |
| Nb24 mammalian supernatant (+ cntrl) | | | |
| Cell-free No DNA (- cntrl) | | | |

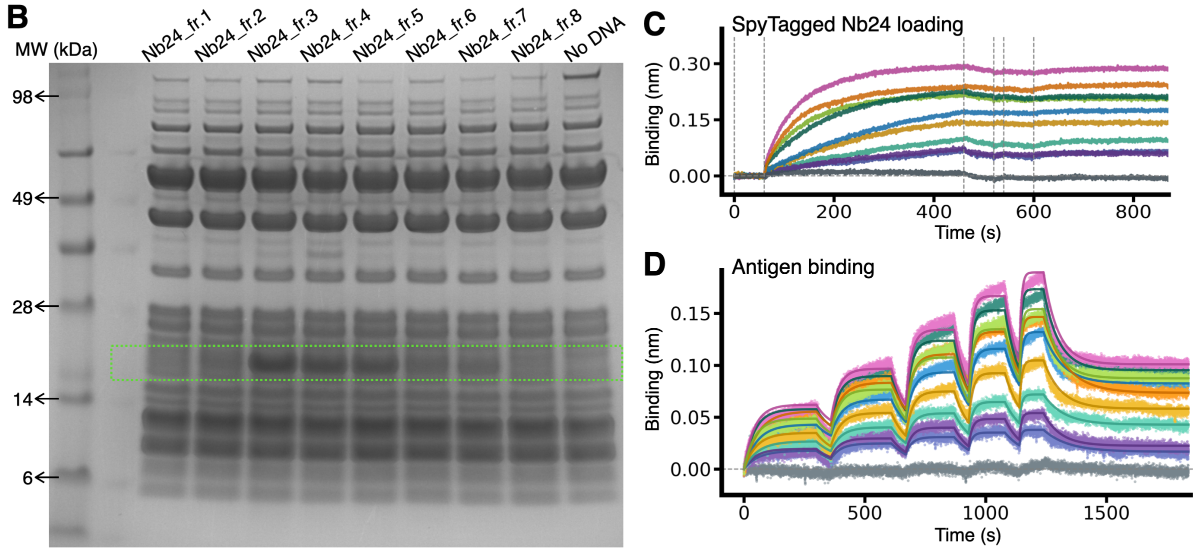


**Figure S3. Optimisation of linear gene fragments for cell-free expression.** (**A**) Table with the tested fragment IDs and their different spacer lengths. Colours correspond to the lines in the plots. Columns are the length of the 5' end of the fragment, the length of the region, which includes the ribosome-binding site, between the T7 promoter and the start codon (ATG), and the presence or absence of a T7 terminator. All DNA sequences are provided in **Supplementary Dataset 1**. (**B**) SDS-PAGE with Instant Blue staining of the cell-free reactions after overnight expression from the different gene fragments (different wells), including a No DNA control, which is used to show the various component of the cell-free expression mixture when no protein is being expressed. An extra band is visible in some wells at just above 17 kDa (inside the green box, which serves as a guide for the eye) corresponding to nanobody Nb24 with a SpyTag003 and a 6xHis tag at its C-terminus (as in **Fig. S1**). The band is most apparent for fragments 3, 4, and 5. (**C**) BLI sensorgrams of the loading of the SpyTagged product to the biotinylated SpyCatcher003 already on the sensor. Sensorgrams have been aligned to the previous buffer step to highlight differences in loading rates, which correspond to differences in expression levels. The grey trace is the negative control showing that no loading is observed from the cell-free expression mixture not expressing a SpyTagged nanobody. The orange trace serves as a positive control and corresponds to SpyTagged Nb24 expressed in mammalian cells and loaded from the cells' supernatant. In agreement with the SDS-PAGE, Nb24 from fragments 3, 4 and 5 loads rather quickly and reaches plateau in less than 400 seconds, while Nb24 from all other fragments loads more slowly, denoting substantially lower expression levels. As expected, no loading is observed from the negative control. (**D**) BLI sensorgrams of the antigen binding steps, aligned to the assay baseline and fitted to a 1:1 partial dissociation binding model (solid lines, except for the negative control). The antigen – $\beta_2$-microglobulin, used as analyte – was present at increasing concentrations of 25, 50, 100, 200, 400 nM in each association phase.

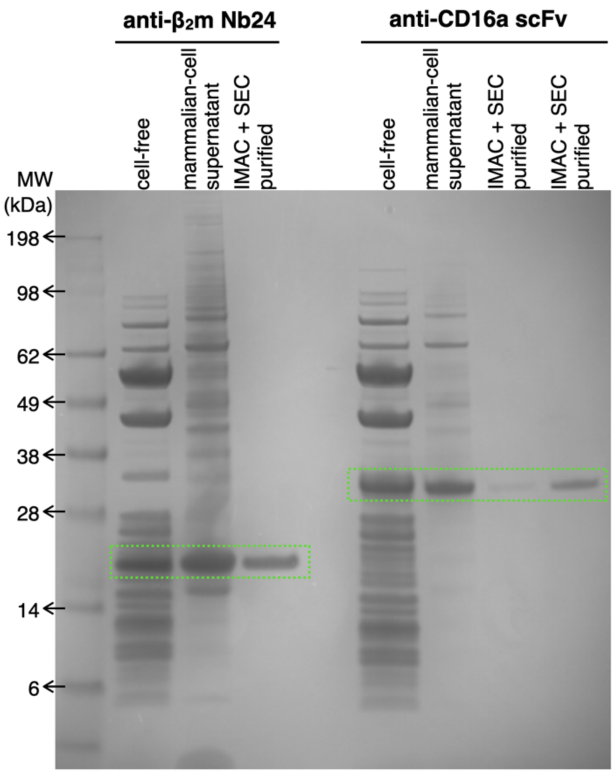


**Figure S4. SDS-PAGE analysis of the binder samples used for generating the data in Figure 2.** Each well corresponds to one of the samples (see header) that were loaded onto the sensor in the experiments presented in **Figure 2** of the main text. The gel was stained with Instant Blue, and the green boxes serve as a guide for the eyes to highlight the bands corresponding to SpyTagged and HisTagged Nb24 or anti-CD16a scFv at their expected molecular weights (see ladder).

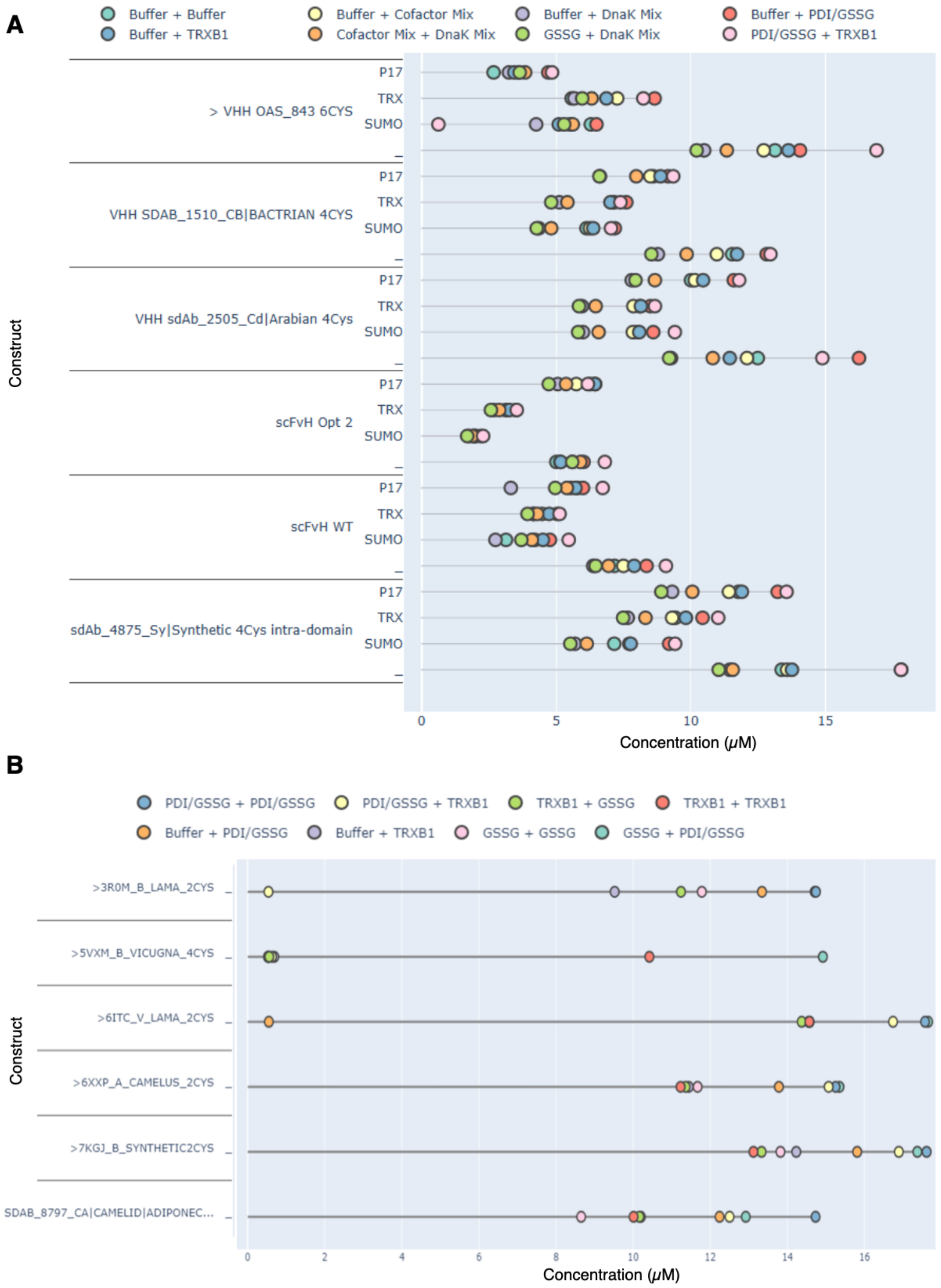


**Figure S5. eProtein Discovery expression conditions screening for nanobodies and scFvs in cell-free blends**. Results are reported as directly visualized from the eProtein Discovery software following an automated 24-hour expression and purification screen. (**A**) Lollipop graph depicting soluble expression yields across 32 combinations of expression conditions and solubility tags, for 6 different binders (y-labels). These conditions were generated from 24 DNA constructs (eGenes), encoding each the 6 binders with three different solubility tags or without any tag (y-label), and tested against eight different cell-free blends as detailed in the legend. There, PDI: Protein disulfide isomerase to promote correct disulfide bond formation GSSG: Mimics the oxidizing conditions of the eukaryotic endoplasmic reticulum and prokaryotic periplasm to promote disulfide bond formation TRXB1: Chaperone to promote correct folding and protein stabilization. (**B**) Like **A** but for six additional nanobodies without any solubility tag and tested against different cell-free blends focussing on oxidising conditions (legend). The data points at around x=0 for the top three nanobodies in the plot were labelled as unreliable by the eDiscovery analysis software, due to droplet collapse on the chip, which most likely resulted from the partial loading of this chip (where we loaded 6 rather than 24 DNA constructs). All other data points were correctly analysed, indicating that the blend with double PDI-GSSG (PDI-GSSG+PDI-GSSG) typically yields the highest expression levels (which is comparable to that from the very similar GSSG+PDI-GSSG blend). The sequences of all constructs are provided in **Supplementary Dataset 1**.

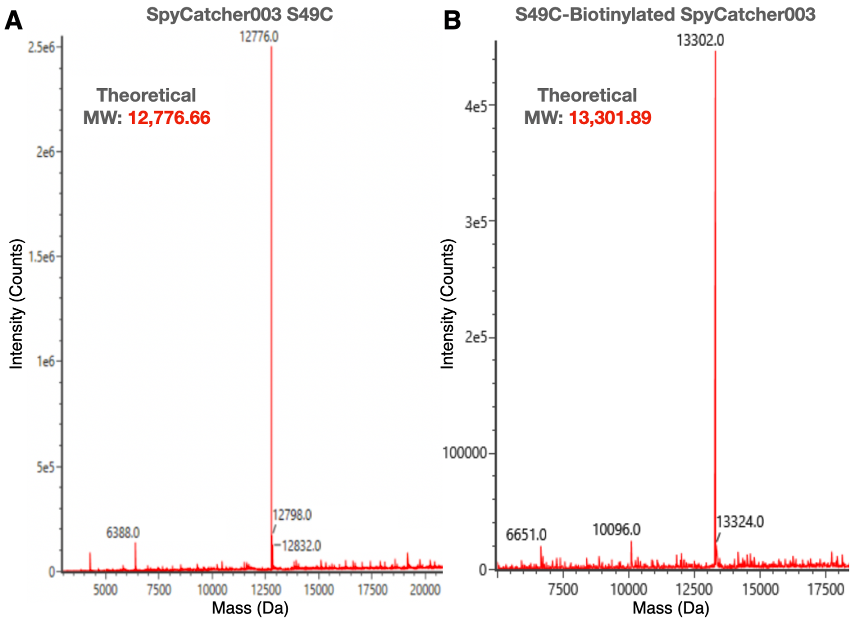


**Figure S6**. Deconvoluted liquid-chromatography mass spectrometry spectra (electrospray ionization) of purified SpyCatcher003 S49C following TEV cleavage (**A**), and of the same protein after maleimide biotinylation (**B**). Expected, theoretical molecular weights (MW) calculated from the amino acid sequence and the maleimide-PEG2-biotin (525.23 Da) are also reported on the panels.

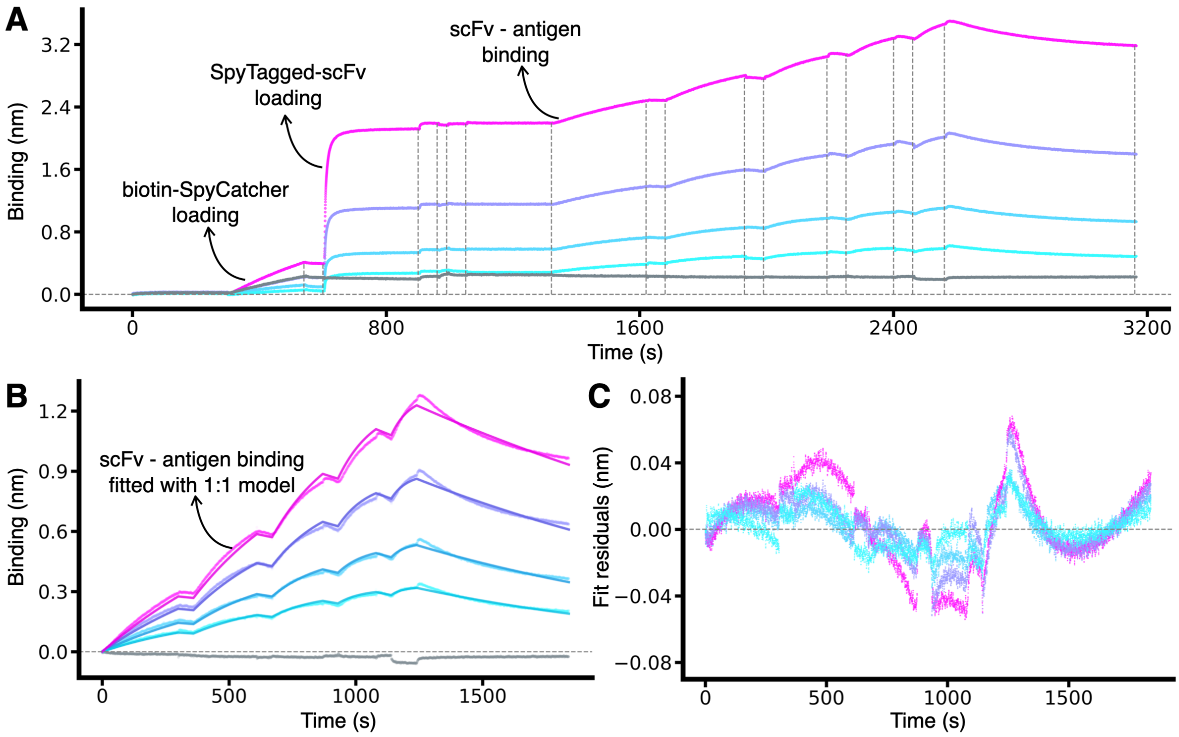


**D**

| SpyCatcher003 Concentration (nM) | Sum of Squared Residuals (nm²) | $K_D$ (nM) |
|---|---|---|
| 12.5 | 0.47 | 4.2 |
| 25 | 0.95 | 4.0 |
| 50 | 2.28 | 3.7 |
| 100 | 4.43 | 3.4 |

**Figure S7. Optimisation of sensor crowdedness for optimal signal quality**. In BLI, the more ligand is loaded the higher the signal to noise ratio is obtained. However, excessive loading leads to large deviations from the theoretical binding models used to fit the data, due to artifacts resulting from surface heterogeneity or mass transport. (**A**) BLI sensorgrams resulting from the loading of different concentration of biotinylated SpyCatcher003 (see legend in **D**) on the streptavidin sensors, and then probing the same wells as detailed in **Fig. 1** of the main text. In all panels, the ligand is always the same anti-CD16a scFv loaded from mammalian-cell supernatant, and the analyte is always the same concentration series of monovalent CD16a (increasing concentrations of 6.25, 12.5, 25, 50, 100 nM). The control for non-specific binding of the analyte (grey trace) was loaded with 50 nM SpyCatcher003, but then not loaded with any SpyTagged protein but blocked with free SpyTag003 peptide in the blocking step (see **Fig. 1**). (**B**) Fits carried out with a 1:1 standard binding model (solid lines) of the analyte associations and dissociations steps. (**C**) Residuals plot of the fits in **B**. (**D**) Table with the different SpyCatcher003 concentrations used at the loading steps, the sum of the squared residuals for the fits, and the resulting $K_D$. The data show that, while all traces yield very similar $K_D$ well within the reproducibility of the technique, the trace obtained with the lowest SpyCatcher003 concentration of 12.5 nM has the best agreement with the theoretical binding model, as shown by the residual plot and the sum of squared residuals.
